# OptiMol : Optimization of binding affinities in chemical space for drug discovery

**DOI:** 10.1101/2020.05.23.112201

**Authors:** Jacques Boitreaud, Carlos Oliver, Vincent Mallet, Jerome Waldispühl

## Abstract

Ligand-based drug design has recently benefited from the boost of deep generative models. These models enable extensive explorations of the chemical space, and provide a platform for molecular optimization. However, current state of the art methods do not leverage the structure of the target, which is known to play a key role in the interaction.

We propose an optimization pipeline that leverages complementary structure-based and ligand-based methods. Instead of performing docking on a fixed drug bank, we iteratively select promising compounds in the whole chemical space using a ligand-centered generative model. Molecular docking is then used as an oracle to guide compound optimization. This allows to iteratively generate leads that better fit the target structure, in a closed optimization loop, without prior knowledge about bio-actives. For this purpose, we introduce a new graph to selfies VAE which benefits from a seventeen times faster decoding than graph to graph methods while being competitive with the state of the art. We then successfully optimize the generation of molecules towards high docking scores, enabling a ten-fold augmentation of high-scoring compounds found with a fixed computational budget.

**Availability:** Code is available on GitHub

## 1 Introduction

### Molecular optimization

Molecular optimization, also known as inverse design, consists in designing compounds that have desired drug-like properties and biological activity. Direct molecular optimization was first introduced in [1]. Subsequently, a string of papers addressed this problem with a variety of approaches as explained in reviews [2, 3]. Formally we are looking for compounds **x** that maximize a function **f**(**x**). This function is often a chemical property that makes a compound more drug-like such as QED or solubility, or a more complex property, such as bio-activity. Several properties of **f** affect its optimization: differentiability, dimension of the output space, evaluation cost and smoothness. Molecular optimization can be further subdivided into lead discovery, which corresponds to this setting, and lead optimization, where we start from a given seed compound, meaning **x** is constrained to a region of the chemical space.

The drug-like chemical space contains an estimated ∼ 10^60^ compounds [4]. The size of the chemical space and the difficulty to accurately estimate the objective **f** without *in-vitro* tests makes it unreasonable to look for just one candidate. Moreover, **f** is often a simple surrogate for a complex, phenotypical endpoint. Formally, we are not after the global maxima of **f**. Hence the current approach to molecular optimization is to search for ensembles of compounds with enhanced estimated properties and then conduct in-vitro tests. We wish to find a non-trivial distribution *q* from which we can sample and that augments 𝔼_**x∼***q*_(**f** (**x**)). The optimal solution would be a trivial distribution always returning the global maxima. To avoid this, we could simultaneously maximize the distribution entropy but in practice, the distribution found is sub-optimal, making it non trivial.

### Binding affinity estimation

In drug discovery, compounds are expected to bind to a given target that can be a protein molecule, an RNA molecule or bigger complexes. To this end, it is crucial to study the binding mode of small compounds and to simulate their binding affinity. There are three avenues for obtaining binding affinity estimates :

- Experimental bio-assays consist in in-vitro quantitative assessment of the interaction between a compound and a target. This does not require computing and is reliable, but often scarce.
- Quantitative Structure Activity Relationship (QSAR) models are machine learning models trained on experimental bio-assays to derive structure-activity rules. They generalize quite well on some targets but as any other machine learning algorithm, they have a validity domain. This means that their accuracy for a compound **x** depends on the similarity between their training set and **x**
- Molecular docking softwares search for ligand conformations that minimize the binding energy with a given protein pocket. Although the estimates of the binding affinity they provide are noisy, the top-scoring compounds are enriched in active molecules [5]. Thus they are widely used to select the most promising leads in a library. Docking is computationally very intensive (∼10 CPU minutes / compound). The small proportion of molecules for which the docking gives insightful results, added to the time cost of docking computations, make it crucial to carefully choose the compounds to dock.

### Binding affinity optimization

In practice, the earliest molecular optimization task consists in finding compounds with high affinity to a given target. In the structure-based approach, an ensemble of lead candidates is obtained by screening fixed libraries using docking. Computational resources currently limit the library size to ∼10^6^ −10^9^ compounds, which is a fraction of the drug-like chemical space.

In the ligand-based approach, QSAR models are used as a surrogate for binding affinity. This approach only implicitly leverages the target structure, and inherently limits the diversity of generated molecules. Therefore, it is not well suited for finding compounds with new binding modes to the target. It also relies on the assumption that molecules with similar structure are likely to exhibit similar bio-activity, which does not hold true in the whole chemical space, especially near activity cliffs [6].

Active learning and automated synthesis have been identified as promising research directions to accelerate the drug discovery process [7, 8]. By allowing a model to iteratively query the most informative data, and learn from experimental answers to these queries, these methods enable a guided and data-efficient exploration of the activity landscape in the chemical space. Inspired by such closed-loop strategies, we propose to iteratively search the chemical space for promising leads, guided by a structure-based assessment of their activity (Figure 1). By combining molecular docking and a latent-variable generative model, our framework finds regions of high affinity to a target in the chemical space.

**Figure 1:**
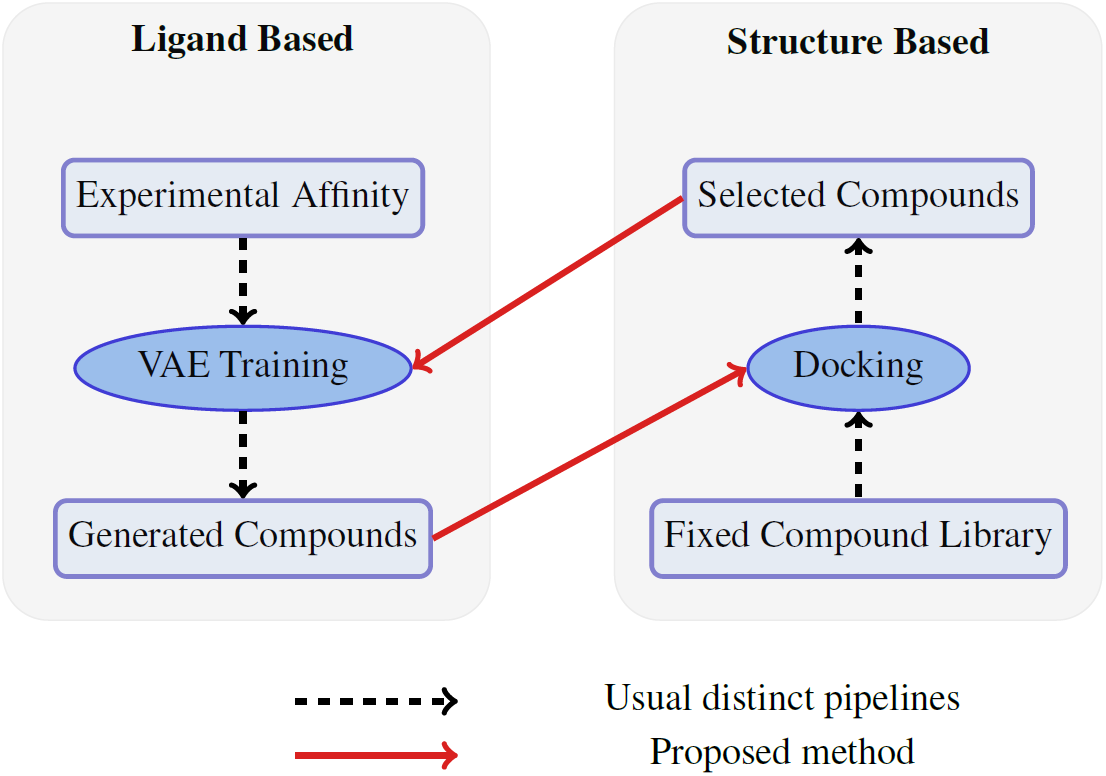
Closing the loop : In ligand-based approaches, a generative model is biased with experimental affinities to produce more active compounds. In structure based approaches, a fixed library is screened for actives. We propose to dock the compounds produced by the generative model, and to use the results of the docking to fine-tune the ligand-based generative model

## 2 Related work

### 2.1 Molecule representation

Molecules can be represented in several ways, trading off between the accuracy of the depiction and its computational advantages. In decreasing order of richness of representation, molecules are represented as ensembles of 3D, static 3D, molecular graphs, SMILES, Selfies [9] or fingerprints. Each of these representation has drawbacks. Using 3D objects with no preferred orientation in machine learning pipelines is under research with promising results [10, 11] but not yet established. Molecular graphs are efficient to encode information but deconvolution and generation is harder [12, 13]. Finally string representations were used first as they benefited from advances in natural language processing. SMILES were extensively used but when generating SMILES, some sequences are invalid. This problem was recently solved by the introduction of Selfies[9] that are all valid sequences, but are sensitive to small modifications.

The choice of the representation of molecules has a direct impact on the optimization process : The discrete nature of sequences and graphs does not allow for continuous optimization. This can be addressed by considering the sampling of the chemical space as a sequential process that can be formulated as a reinforcement learning problem [14]. Another approach uses Variational Autoencoders (VAE) [15]. These methods learn a continuous chemical space by reconstructing or translating these representations [1, 9, 12, 16, 17, 18, 19]. The latent space can be interpreted as a flattened manifold, of which the structure and geometry mostly reflect chemical similarity. Turning the chemical space into a euclidean space enables to continuously navigate in the space, which opens the door to classical optimization methods.

### 2.2 Molecular optimization

#### Latent space optimization

Once molecules are embedded in a continuous space, classical optimization schemes become possible. Bayesian optimization was used first [1, 12, 20, 21], followed by constrained Bayesian optimization [22, 23] and swarm optimization [8]. Another approach is to approximate **f** with a differentiable function and to conduct gradient ascent in the latent space. A potential advantage of this method is that the function mapping from latent space to **f** can be learned jointly with the generative model, contributing to shape the latent space [17].

#### Molecular translation

Matching Molecular Pairs Analysis were recently used by [24] to train a translational VAE to turn a molecule into a similar one with better target property: this model learns to take an optimization step, given a starting point. In [25], the authors show that molecules can be optimized for a target property by recursively taking such steps in the chemical space. This step-wise approach is limited to lead optimization, but seems well suited to handle activity cliffs.

#### Guided generation using Reinforcement Learning

By decomposing the molecule generation as a sequence of actions, we can build a generative model using a probabilistic RL agent. Guided generation consists in biasing the generation towards compounds that optimize **f**, easily including non differentiable objectives [14, 26]. An adversarial term can be added to ensure that we keep generating realistic compounds [27, 28]. One caveat in using these methods is that for multidimensional objective functions, the rewards get sparse and the training gets harder [27]. Another caveat is that the number of evaluation of **f** is not optimized, limiting the use of costly oracles.

#### Other generative models

The authors of [29] use a fragment based approach where they automatically learn a library of activity-inducing fragments and then generate compounds combining them. In [30], a Generative Adversarial Network (GAN) is trained to mimic the active compounds distribution in the latent space. A related approach was proposed in [31], and performs iterative fine-tuning of a generative model on its most successful outputs. Other approaches of this kind with better statistical grounding exist [32, 33], but we are unaware of their application to small molecule optimization. They are applicable both to models with and without a latent space structure. In [34], the authors condition a generative model by a 3D shape. By coupling it with a GAN that generates shapes conditioned on a target protein structure, [35] generates ligands that fit an input protein structure. This a way to generate compounds with augmented affinities that side-steps the direct optimization.

### 2.3 Binding affinity optimization

To generate compounds with high binding affinities, we can also use one of the three aforementioned sources of binding affinity estimates. In [30], authors train a GAN to sample from a distribution resembling the one of known actives in latent space, and evaluate the samples affinities using a QSAR model. Alternatively, a QSAR model can be used to bias the generation. This is the approach used in the RL framework by [14, 26]. This method gives good results, but is based on a QSAR model that can be inaccurate: the authors constrain the model to avoid drifting away from its validity region. We circumvent this limitation by resorting to a docking program, that uses physics-based molecular mechanics force fields to compute binding affinities.

### 2.4 Contributions

In this work we introduce a new VAE that is more computationally efficient while retaining state of the art results. We then turn to binding affinity optimization using this generative model. We use docking as an oracle and include docking score in the function **f**, making it non differentiable. In addition, this oracle is now costly and the sparse rewards induced by the RL frameworks are not tractable, which calls for specific optimization methods. Bayesian Optimization (BO) is well established and optimizes queries, but has limitations in high dimensions and does not yield a generative model. We explore the use of the recently published Conditioning by Adaptive Sampling (CbAS) [33].

## 3 Methods

### 3.1 Graph to sequence Variational Autoencoder

Inspired by previous Variational Autoencoders architectures, we propose a graph to sequence VAE that achieves comparable performance to state of the art models, while benefiting from the following design choices :

- Using the molecular graph as input and a graph convolution encoder solves the issue of data augmentation. It is also better suited for learning a chemically organized latent space, since chemically similar molecules have very similar graphs, while their SMILES representations may change more due to syntax rules. Finally, graph convolution embeddings and circular fingerprints were shown to enhance molecular properties predictions [36, 37].
- On the decoding side, however, decoding to a molecular graph results in more complex architectures than decoding to sequences, thereby increasing the computation time. The main weakness of string-based decoders is that due to the sensitivity of the SMILES syntax, only a small fraction of the decoded sequences resulted in a valid molecule. By using recently published Selfies [9], we circumvent this issue and generate 100% valid molecules.

We train our model on the Moses benchmark set [38], which contains ∼1.5M molecules from the ZINC database [39] clean leads. The model architecture and training regime are detailed in A.1.

### 3.2 Docking on Dopamine Receptor D3

We use Autodock Vina [40] to estimate compounds binding affinities to human Dopamine Receptor D3. We use the PDB structure of the DRD3 receptor provided in the DUD-E data set [41], and keep the same binding site coordinates. We set the exhaustiveness of the conformation space search to 16, as a reasonable compromise between running time and enrichment factor. We compute the docking score as the average of the 10 best poses found for each ligand.

### 3.3 Lead Generation

To generate promising leads for a target, we wish to sample in the chemical space from a distribution *q*(**z**) that maximizes the docking score oracle **f**, while generating a diversity of molecules. Reinforcement learning methods require many evaluations of **f**, and therefore seem ill-suited. As **f** is not differentiable, we also exclude gradient-based methods.

#### Bayesian Optimization

uses Gaussian processes to approximate **f** in a query-efficient way, by learning on points that maximize expected improvement. To sample these points, a rigid (not learnt nor adaptive) sampling is used. This does not scale well to a high dimensional space and for a large batch size as the grid evaluation becomes intractable (This amounts to finding an estimate on all molecules and only picking the most promising candidates for docking). This choice limits scalability when sampling tens of thousands of compounds in the chemical space.

#### Conditioning by Adaptive Sampling

is a recently published method [33] that addresses the limitations of Bayesian Optimization. This method trains a generative model that also seeks to maximize an objective function. Starting from a prior generative model, it progressively shifts its distribution to maximize the expectation of a function of the samples. Queries are used in an efficient way thanks to an importance sampling scheme coupled with reachable objectives for the model. The alternating phases of tuning and sampling also enable a computationally efficient implementation. For a more detailed explanation of these algorithms, see A.3 In the specific case of affinity optimization, as a prior model, one can either use a model trained on a broad chemical space, or leverage previously discovered actives to narrow-down the chemical space by fine-tuning the prior on the actives. Then a docking program is used to re-weight the samples and fine-tune the model.

#### Efficient implementation

To our knowledge, this is the first use of CbAS with a costly oracle. We already mentioned the computational cost of docking, so using CbAS required a dedicated implementation, alternating between a phase of sampling and re-weighting compounds, a phase of docking and a phase of model training. We have implemented a parallel version of the code that can leverage a multi-node architecture cluster for the docking phase using several hundreds of CPU cores and running the sampling and training phases on GPU nodes. This implementation made the training run 30 iterations over batches of 1000 samples in approximately 15 hours using 200 cores. We note that parallelism over more CPU would result in linear speed up, up to several thousands of CPU. The implementation is available on GitHub and easily adapts to any pocket structure.

## 4 Results

### 4.1 Graph to Selfies VAE

Moses[38] is the reference work for benchmarking molecular generative models. Moses defines a standard train/test split of a given data set and a standard way of computing metrics. Our model achieves state of the art performances while benefiting from design choices in our application. Noteworthy, our model has 3.2M trainable parameters, compared to 5.1M for the default JTVAE implementation [12]. Our model also runs approximately 17 times faster for sampling and training. We achieve comparable results on the Moses benchmark metrics (Table 1, see Supplementary Table A.1 for full comparison).

**Table 1:**
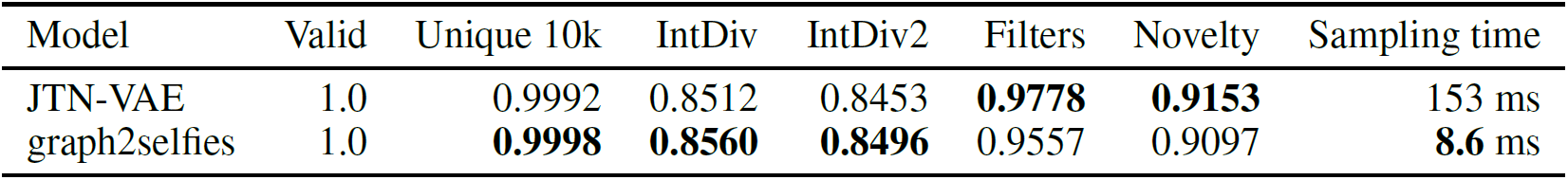
*Left* : Moses metrics for samples generated by JTVAE and graph2selfies. *Right* : Sampling time is the average time needed to produce one molecule from a trained model.

To check our latent space is suitable for optimization tasks, we run Bayesian Optimization on the toy task of enhancing the composite logP score^1^, and compare to latent spaces introduced in previous works. As all baselines, we use the open-source implementation in [20] and report our results on Table 3. We note that picking only the 3 best was subject to high variance as we witness changes in performance when training the gaussian process differently. We conclude that our architecture has a similar performance as the current state of the art with a significantly reduced computational load.

**Table 2:** Enrichment Factor (EF) in active compounds at different thresholds

| EF | Threshold |
| --- | --- |
| 13.373 | 0.1% |
| 11.122 | 0.5% |
| 8.168 | 1% |
| 7.229 | 2% |
| 8.919 | 1% decoys |

**Table 3:** Top 3 scores found by each method (All scores are normalized using the 250k dataset scores, base-line results are copied from previous works).

| Model | 1st | 2nd | 3rd |
| --- | --- | --- | --- |
| GVAE [20] | 2.94 | 2.89 | 2.8 |
| SD-VAE [21] | 4.04 | 3.5 | 2.96 |
| G2S - BO | 4.81 | 4.6 | 4.27 |
| JTVAE [12] | 5.3 | 4.93 | 4.49 |
| GCPN [27] | 7.98 | 7.85 | 7.8 |
| G2S - CbAS | **10.88** | **10.57** | **10.5** |

**Table 4:** Mean pairwise Tanimoto distance in 4000 samples produced by sampling from different distributions

|  | Tanimoto distance |
| --- | --- |
| Moses samples | 0.89 |
| CbAS samples | 0.87 |
| ZINC samples | 0.90 |
| Skalic *et al.*[35] samples | 0.89 |

### 4.2 Docking on Human Dopamine Receptor 3

To evaluate the ability of docking to find actives for DRD3, we dock 380 clustered actives and 20 000 decoys from the ExCAPE database [42]. Full docking scores distributions are shown in Supplementary Figure A.2. The log-ROC curve (Figure 2) shows the fraction of actives found at a given decoys fraction threshold, with a focus on the top-scoring compounds in the ranked list. When compared to a random permutation of the list, it shows that docking manages to separate actives and inactives, with significantly more actives in the top of the list.

**Figure 2:**
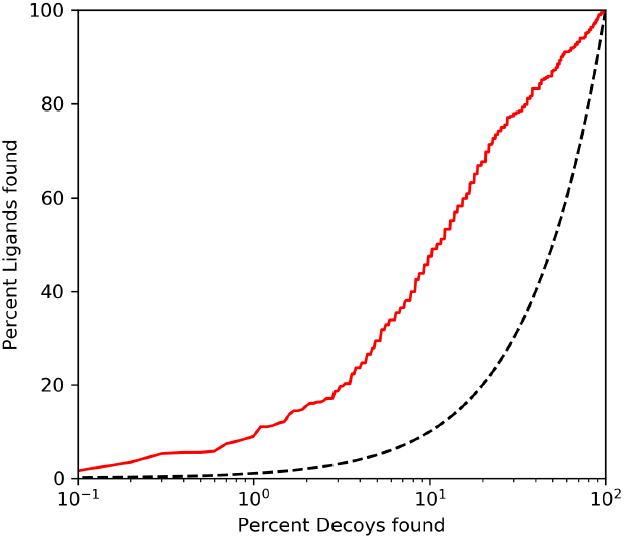
*(red)* Log ROC curve for ExCAPE database DRD3 actives and inactives *(dashed line)* Random sampling in the ligands and decoys set

We then compute enrichment factors at different thresholds (Table 2). Among the 1% top-scoring compounds, we find a ratio of actives 8.168 times higher than in the whole list. In [41], the authors compute such metrics for 102 diverse proteins. It is worth noting that they obtain an enrichment factor at 1% decoys (EF1) of 4 for DRD3, which is quite low compared to other targets, as several get an EF1 higher than 40. This suggests that DRD3 is a rather difficult target for docking softwares, but the docking oracle we use still provides useful information to guide the generation towards DRD3 actives.

We conclude that our docking oracle is able to enrich a list of samples in actives. Hence, the top-scoring compounds found by docking are significantly more likely to bind DRD3. As a control for lead generation experiments, we also dock 10% of the Moses training compounds [38]. As expected, their docking scores distribution is significantly shifted from the actives distribution, and only a negligible fraction of them have a score better than −10 kcal.mol^−1^ (Supplementary Figure A.2).

### 4.3 Optimization using CbAS

To compare with previous optimization methods, we first apply CbAS to the toy task of enhancing the composite logP (clogP) score. This is the same benchmark as in Table 3. We obtain scores (normalized, as in Table 3) of 10.88, 10.57 and 10.5 respectively. CbAS outperforms the Reinforcement learning approach in [27]. In Figure 3, we plot the evolution of the cLogP scores distributions during the optimization process, as we find this metric more robust than the best samples. Top-scoring compounds are showed in Supplementary Figure A.3.

**Figure 3:**
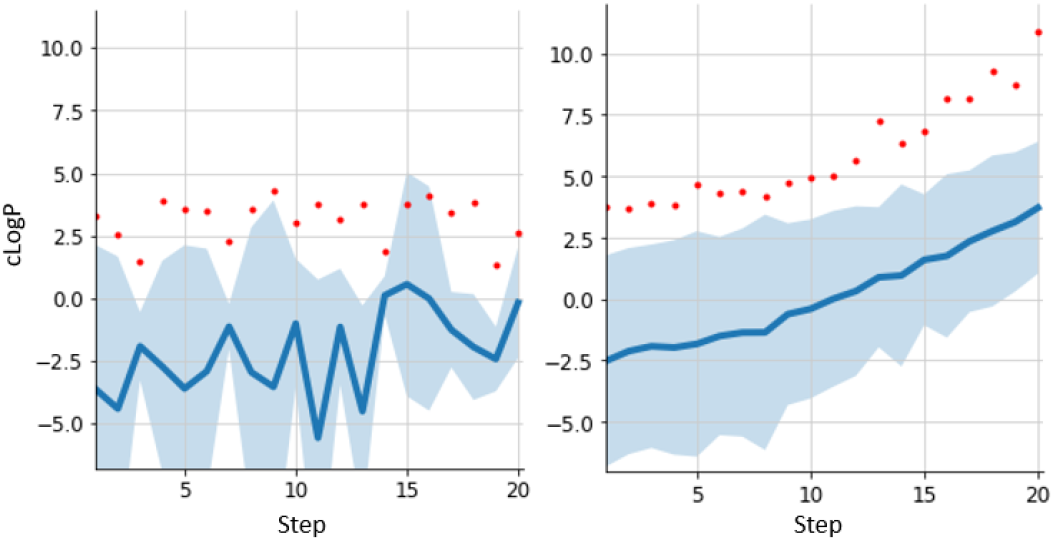
*left:* Bayesian Optimization of clogP with 10k initial samples and 500 new samples per step, using implementation from [20]. *right:* CbAS for clogP optimization, with 1k samples per step

#### Query efficiency

As stated earlier, query-efficient methods are necessary for affinity optimization. This is due to the computational cost of docking, our end goal. The methods in Table 3 are not query efficient because the queries in a RL model are not optimized to be maximally informative and that there is no alternated phase of sampling and training. The implementation of BO from [20] used as a benchmark earlier uses annotation of the whole 250k compounds for training. This is not a reasonable setting for real-life applications, as it is unlikely that all training compounds are labeled. To compare with CbAS, we use another implementation of BO that uses less data for training (Supplementary A.3). This implementation is still slower than CbAS and does not support batches of 1000 samples. In addition, this method would definitely not be suitable for extensive sampling of tens of thousands of compounds once trained. Still, it is the only other query efficient method in this benchmark. Details about settings and implementation are in Supplementary A.3. We compare this method to the run with CbAS.

As can be seen in Figure 3, CbAS has a smooth improvement over epochs. It manages to find compounds with a higher and higher cLogP, above any other methods. If we optimize for too long, we start having unrealistic samples leveraging the weakness of logP computational estimation. This was also found to happen in the runner up GCPN and is illustrated in Figure A.3. The trade-off between similarity to prior molecules and clogP enhancement can be controlled by early stopping in the CbAS training. We see that we can get reasonable compounds after 15 epochs, with better cLogP scores than the state of the art, showing the efficiency of our approach.

### 4.4 Docking optimization

We now come to the main result of our work : The generative model learned using CbAS generates samples with enhanced docking score (shifting the average score from −7.5 to −8.5 kcal.mol^−1^) while preserving chemical diversity as is shown in Figure 4. This means that we managed to generate a population with a docking score distribution close to the actives one, while being able to sample diverse compounds. As a prior, We used the model trained on Moses as it contains clean leads that are suitable for docking optimization.

**Figure 4:**
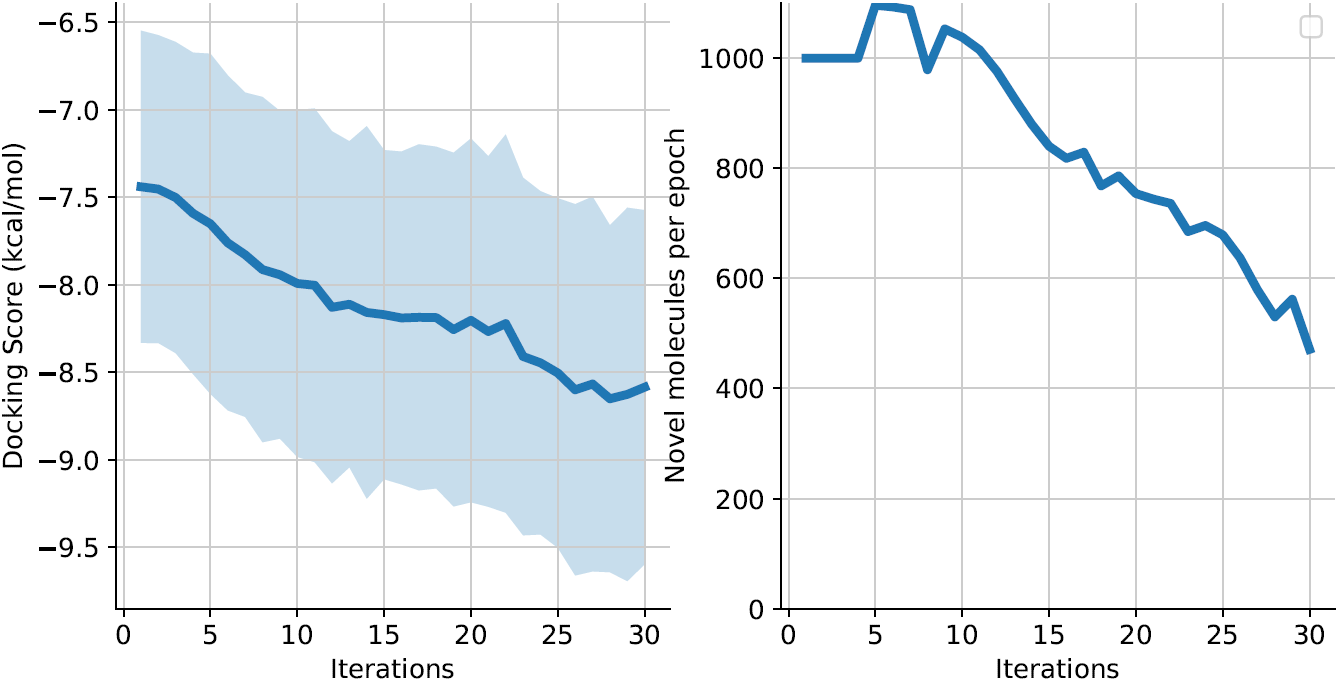
*Left*: The average and standard deviation of the docking scores obtained at each CbAS iteration. *Right*: The number of compounds never sampled before, sampled at each CbAS iteration.

We compare to [35] that also try to generate actives for a target, and assess their method by docking the sampled compounds. It is worth noting that this comparison is not done in an ideal setting as DRD3 is in their training set and their molecular generative model is trained on the ZINC database [39], but we check that this prior has the same docking scores distribution as MOSES (Fig 5). We note also their model is a different pipeline that can take as input any target without the need for retraining a generative model, as a shape captioning network governs the sampling. Another fundamental difference with CbAS is that We can keep fine-tuning the model further and further to get better scores, at the cost of loosing the diversity contained in our prior model. This is not possible with [35], where the distribution of the samples is determined by the shape prototypes extracted from the target. To the best of our knowledge, this is the only approach that manages to enhance docking scores. We now further evaluate the generative model obtained through our procedure.

**Figure 5:**
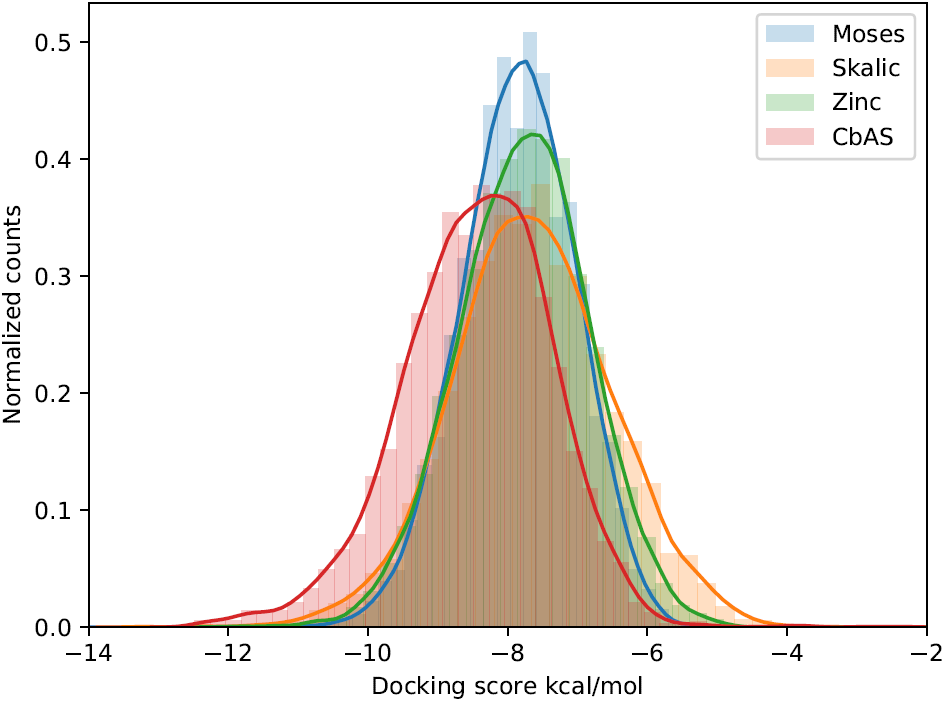
Distributions of scores of docking for 10% Moses training set, 4k CbAS samples, 10k random compounds from ZINC molecules used as prior in [35] and 4k samples generated with [35]

#### Sampling

We want to assess if we are able to generate a population from the new generative model, and if this population has better affinities than the baseline. We easily sampled 100k unique compounds in about one hour after the model was fine tuned. The distribution of docking score of a sub-sample of 4000 molecules from this sampling is available in Figure 5. We see that the compounds drawn from the generative model have a significantly better score than random samples from Moses, closer to the active ones (respective means and standard deviations −8.5 +/−1.1, −7.5 +/− 0.8 and −9.6 +/− 1.2 kcal.mol^−1^). We also apply the method proposed in [35] to our target. It does not really shift the distribution from its prior (ZINC compounds, Figure 5). Under the ROC-AUC metric that they introduce – ability to distinguish enhanced sampling vs random one -, they get a score of 0.47 while CbAS after 30 iterations gets a score of 0.67. This score could be augmented by pushing further the optimization process, but this would be at the cost of diversity.

We also check that the internal diversity of the compounds did not collapse during optimization. We use the mean pairwise Tanimoto distance as a diversity metric. This distance is a bit-to-bit distance over fingerprints, ranging from zero to one. Table 3 shows the results we got for random samples taken in Moses compared to ones derived from CbAS sampling. We can see that there is no major difference, indicating the samples produced by our model remain chemically diverse.

We have also assessed whether the samples deteriorated during sampling. Indeed to get unique samples, we used rejection sampling making the first samples the most likely under the generative model. We found that the difference between the first and the last of the 100k samples was statistically significant, but relatively small with respective means of −8.6 and −8.4 kcal.mol^−1^ respectively. The respective distributions are available in Supplementary figure A.4. The 50 top-scoring molecules from 4k random samples drawn from the CbAS generative model are shown in Supplementary Figure A.5. We note the presence of macrocycles among them, which may be actives but are not drug-like molecules.

This suggests the use of a composite objective function, to penalize the docking score by drug-likeliness or synthetic accessibility.

These results highlight the potential of our method to speed-up lead discovery without making compromises on the exploration of the chemical space. Indeed, in our experimental setting where Moses is used as the prior chemical space, only 0.62% of the compounds have a docking score better than −10 kcal/mol (estimated on a sample of 10% of Moses). In contrast, at the cost of docking 30k compounds to train the CbAS generative model, we are able to sample from a distribution where 7.4% of the samples have docking scores better than −10. If we were to dock 100k random molecules from the prior chemical space, we therefore would expect to find 620 compounds with a score better than −10. Training the CbAS generative model required 30k docking queries, but the expected number of hits with score better than −10 if we then dock an additional 70k compounds is 5180, almost 10 times higher.

## Conclusion

In this paper, we have introduced a new latent space representation for small molecules using graphs and Selfies and show that we obtain state of the art representation with lower computational cost. Using this model, we introduce a new task of building a generative model to create samples with enhanced docking scores, in a feedback loop manner. We show that we are able to train this model so to generate compounds with better and better docking scores.

This method could benefit ligand-based pipelines by generating compounds with enhanced activity towards a target. This would allow for target optimization without having to rely on previously available affinity data. This is useful for novel targets such as the ones involved with SARS. It could also benefit structure-based pipelines by generating focused libraries for docking and saving computations on low affinity compounds, as is demonstrated by the tenfold increase in the ratio of hits at fixed budget. Finally it can be seen as a step towards using both approaches in a unified drug discovery pipeline.

We plan to extend the validation by retraining of the prior on other data sets. In addition, we hypothesize that the smoothness of the objective function may impact the success of the method. Future work could investigate latent space reshaping, using contrastive methods and molecules 3D properties.

## Acknowledgments

We thank Jean Philippe Vert, Olivier Sperandio, Arnaud Blondel, Guillaume Bouvier and Laura Ortega for helpful feedback and discussions.

## A Supplementary Material

### A.1 Model architecture and training

The encoder consists in 3 Relational-GCN layers of hidden size 32, with skip connections, resulting in 96-dimensional embeddings. Two dense layers map to the mean and log standard deviation of the latent embeddings, of dimension 56. The decoder is a 3-layer GRU with hidden states of dimension 450. The model was implemented in PyTorch [43] and DGL [44]. The model was trained for 50 epochs using Adam optimizer, a learning rate of 10^−3^ and an exponential decay with rate 0.9 every 40k steps. Batch size was set to 64. For the first 40k steps, we train the model only on reconstruction loss, and then progressively increase the weight of the KL term by 0.02 every 2k steps, until it reaches 0.5

**Figure A.1:**
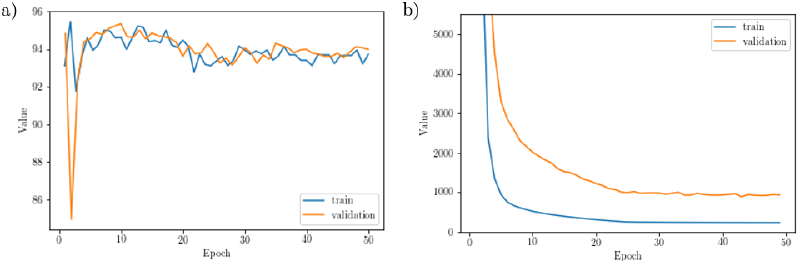
Fraction of correctly reconstructed characters (a) and KL divergence term (b) during model training, for training and validation set

**Table A.1:** Moses metrics for all models benchmarked in Moses[38] and graph2selfies

| Model | Valid | Unique 1k | Unique 10k | IntDiv | IntDiv2 | Filters | Novelty |
| --- | --- | --- | --- | --- | --- | --- | --- |
| HMM | 0.0760 | 0.6230 | 0.5671 | 0.8466 | 0.8104 | 0.9024 | **0.9994** |
| NGram | 0.2376 | 0.9740 | 0.9217 | 0.8738 | 0.8644 | 0.9582 | 0.9694 |
| Combinatorial | 0.9979 | 0.9983 | 0.9948 | 0.8812 | **0.8741** | 0.7912 | 0.9913 |
| CharRNN | 0.9748 | **1.0000** | **1.0000** | 0.8562 | 0.8503 | 0.9943 | 0.8419 |
| AAE | 0.9368 | **1.0000** | 0.9973 | 0.8557 | 0.8499 | 0.9960 | 0.7931 |
| VAE | 0.9767 | **1.0000** | 0.9984 | 0.8558 | 0.8498 | **0.9970** | 0.6949 |
| JTN-VAE | **1.0000** | **1.0000** | 0.9992 | 0.8512 | 0.8453 | 0.9778 | 0.9153 |
| LatentGAN | 0.8970 | **1.0000** | 0.9970 | 0.8565 | 0.8504 | 0.9727 | 0.9488 |
| graph2selfies | **1.0000** | **1.0000** | 0.9998 | 0.8560 | 0.8496 | 0.9557 | 0.9097 |

### A.2 Docking scores distributions

**Figure A.2:**
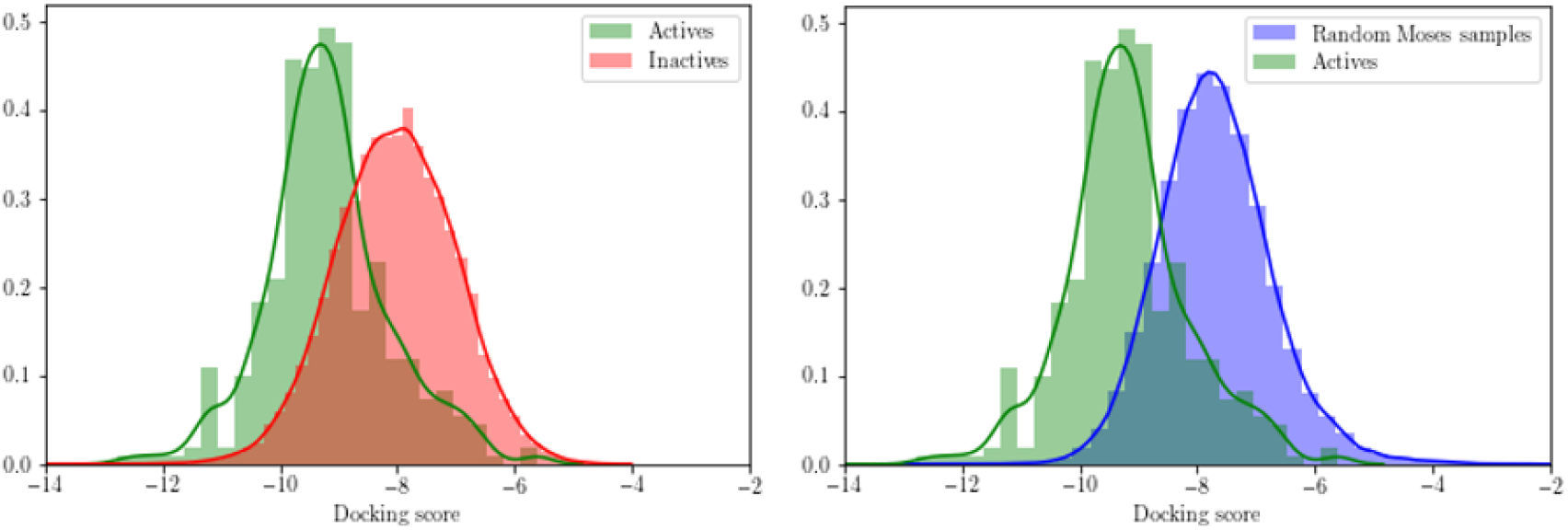
Distribution of docking scores of active molecules vs ExCAPE inactives *(left)* and active molecules vs random Moses compounds *(right)*

### A.3 Bayesian Optimization

Bayesian optimization uses Gaussian processes as a surrogate model for black box functions that are costly to evaluate. It enables to optimize the objective in a query-efficient way, by sampling points that maximize expected improvement. However, it has inherent limitations for the lead generation problem we attempt to solve. To take samples, some kind of rigid (not learnt nor adaptive) sampling is generally used, meaning that the expected improvement under the Gaussian process model is computed over each point of a grid of a certain resolution.

This computation of the expected improvement over the samples space does not scale well to a high number of dimensions for a large batch size as the grid evaluation becomes intractable (This amounts to finding an estimate on all molecules and only picking the most promising candidates for docking). In addition, BO was shown to perform better when the latent space is shaped by the objective function [1, 22]. This is not the case for binding affinities, since the chemical space is likely to exhibit activity cliffs and scattered activity peaks. The final goal is to be able to take samples from the model, and this method uses rejection sampling to generate favorable samples. This choice limits scalability when sampling tens of thousands of compounds.

Bayesian optimization was implemented using BoTorch [45]. A Gaussian process was trained to predict the objective on 500 initial samples selected to be maximally diverse in the Moses training set. The Gaussian process was then trained for 20 steps by sampling a batch of 50 compounds using Expected Improvement as the acquisition function. For the optimization of QSAR activity scores, the 500 initial samples were selected as maximally diverse from the union of Moses and the QSAR train actives.

### A.4 Conditioning by Adaptive Sampling

#### Formulation

To address the limitations of Bayesian Optimization, we turn to a recently published method : Conditioning by Adaptive Sampling (CbAS) [33]. This method trains a generative model that also seeks to maximize an objective function. This methods uses a prior generative model and shifts its distribution to maximize an expectation. Queries are used in an efficient way thanks to an importance sampling scheme coupled with reachable objectives for the model. The alternating phases of tuning and sampling also enable a more efficient implementation.

CbAS starts with a prior generative model with parameters ***θ***^(0)^, *p*(**x** |***θ***^(0)^) and the optimization is formulated by conditioning this probability on the random variable ***S***, *p*(**x**|***S, θ***^(0)^). This random variable represents the values for a probabilistic (or noisy) oracle : ***S*** = (**f** (**x**) *> γ*)|**x**. We now see that this conditioned probability model is a distribution that maximizes the function **f** when *γ* goes to the maximum value of the function. However this conditional probability is intractable and the authors propose to use variational inference to approximate it. The parametric family is a generative model with parameters ***ϕ***, *q*(**x**| ***ϕ***) that solves :

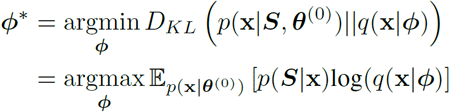

The authors then use importance sampling instead of always sampling from the prior to estimate this expectation. The proposal distributions are the successive generative models obtained at each iteration. The last key idea is to use a fixed quantile of the successive generative models distributions as a value for *γ*^(*t*)^, to set reachable objectives for the model : ***S***^(*t*)^ = **f** (**x**) *> γ*^(*t*)^. A detailed derivation can be found in the original paper and result in solving for ***ϕ***^(*t*)^ for each *t* in the following equation :

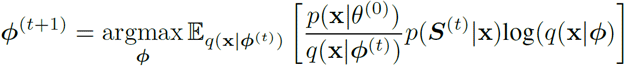

As a prior model *p*(**x** |***θ***^(0)^), one can either use a model trained on a broad chemical space, or leverage previously discovered actives to narrow-down the chemical space by fine tuning the prior on the actives. Then, we take samples from the search model (initialized with the prior), use an oracle with a noise model to get *p*(***S***^(*t*)^ |**x**) and fine tune the search model with the adequately re-weighted samples.

#### Results

The top 3 molecules found with CbAS after 12, 16 and 20 steps are shown in Figure A.3. Like in [27], we note that the best molecules look significantly different from the ZINC clean leads of the 250k dataset. This is due to the nature of composite logP objective, and more reasonable molecules, though lower-scoring, can be obtained using early-stopping with CbAS.

**Figure A.3:**
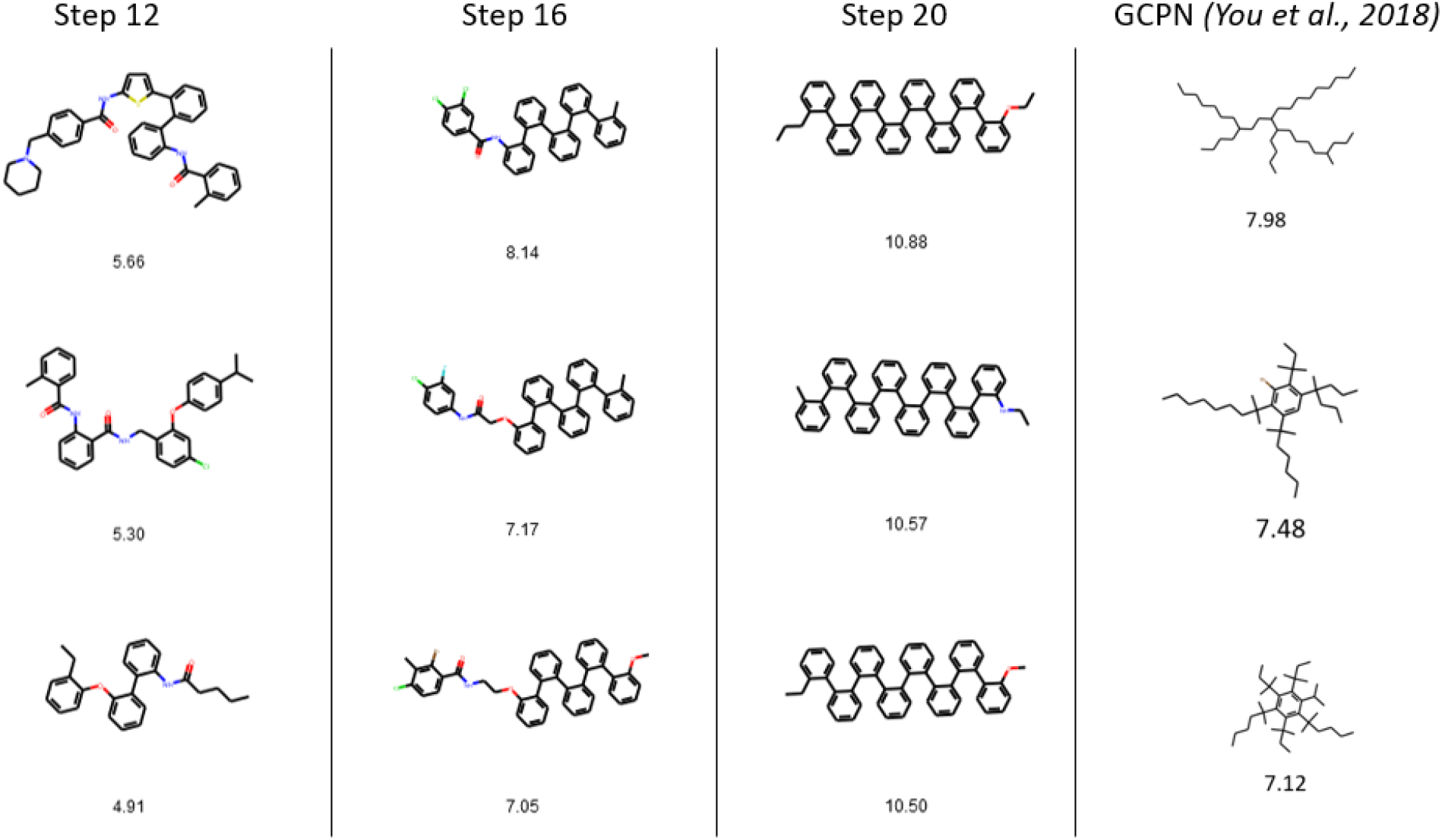
*(left)* Top 3 molecules found with CbAS after 12, 16 and 20 steps *(right)* Top 3 molecules reported by [27]

#### Model optimization

We have tried tuning several parameters and found the training dynamics quite subtle, as too much training increased the scores but crashed the diversity and too little did not optimize the objective functions. We have tuned the number of samples at each epochs, the number of epochs, the usage of teacher forcing, different noise model for the oracle, the number of epochs, the optimizer used (Adam or SGD), the scheduler used, the initial learning rate, the impact of the quantile picked as well as using a non-decreasing *γ*. We found that that the best combination was reached using 30 epochs of 1000 samples, teacher forcing, gaussian noise of variance of the same magnitude as the variance of the data, Adam optimizer with no scheduler and default learning rate, 6th quantile and a non-decreasing *γ*.

#### Sampling quality assessment

Figure A.4 shows the distributions of docking scores for the first new samples from the generative model and the last ones. As stated in the main text, the distributions are statistically significant (p-value of 10^−14^) but small compared to the difference with uniform sampling over Moses.

**Figure A.4:**
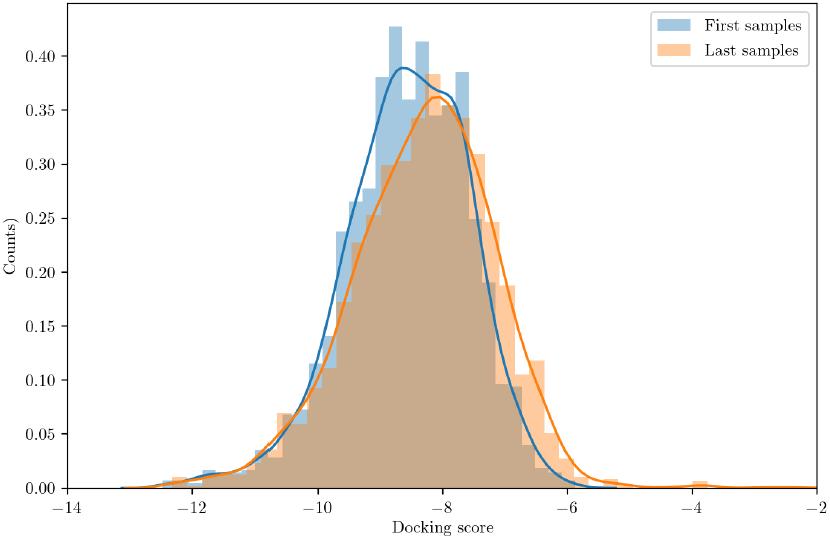
Distributions of docking scores for the first 2000 samples versus the last 2000 samples

**Figure A.5:**
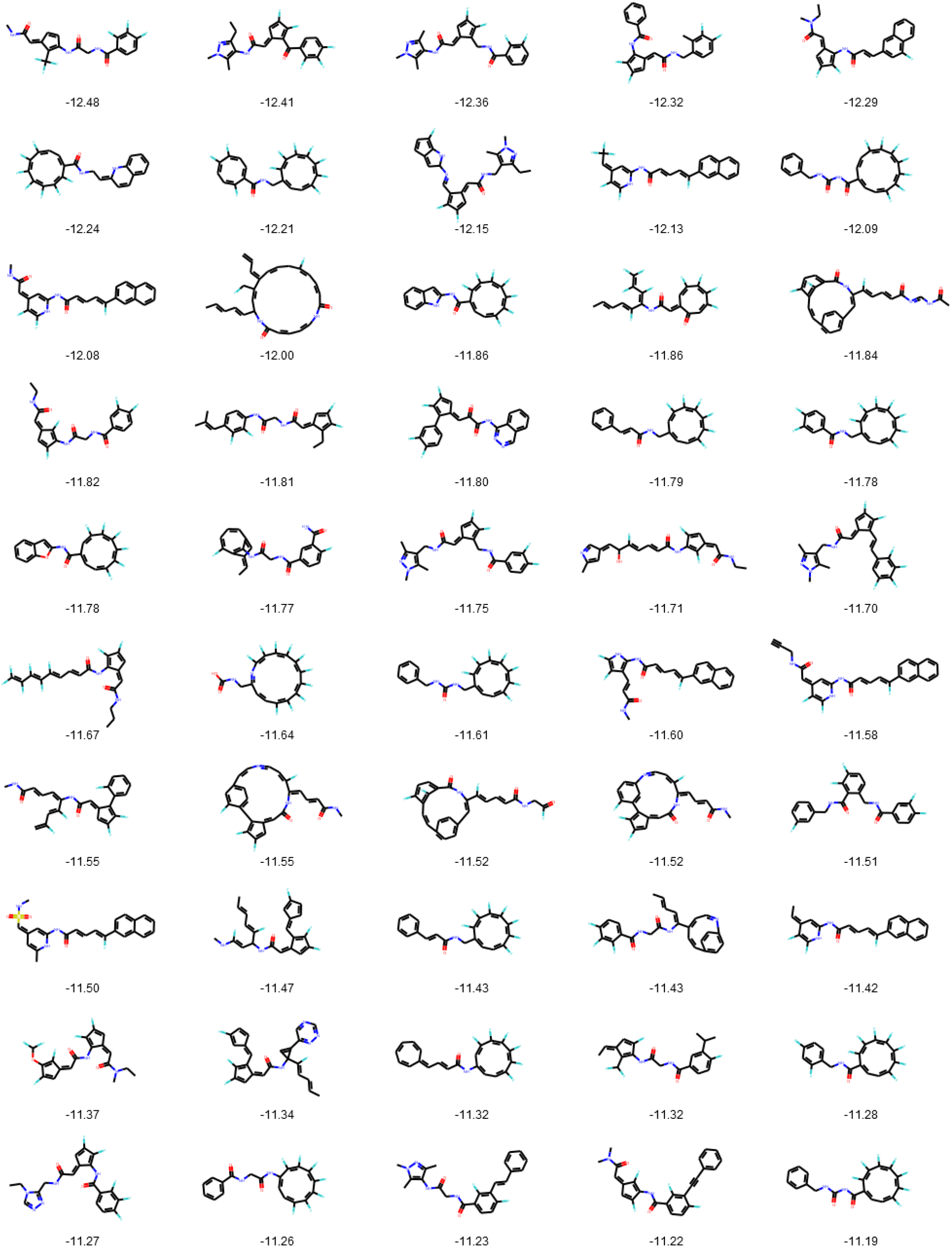
50 best scoring molecules in a random sample of 4000 compounds drawn for the CbAS generative model

## Footnotes

1 *clogP* (*m*) = *logP* (*m*) − *SA*(*m*) − *cycle*(*m*) where *cycle(m)* is 0 if the molecule has no cycle bigger than 6 atoms, otherwise the size difference between the biggest cycle in the molecule and a 6-atoms cycle

